# Functionalized Biomimetic Hydrogels Enhance Salivary Stem/Progenitor Cell Organization

**DOI:** 10.1101/2021.08.05.455302

**Authors:** Mariane Martinez, Robert L. Witt, Mary C. Farach-Carson, Daniel A. Harrington

## Abstract

Complex branched salivary structures remain challenging to replicate within implant ready hydrogels. We showed previously that hyaluronic acid (HA)-based hydrogels enable growth and organization of primary salivary-derived human stem/progenitor cells (hS/PCs) into multicellular spheroids. Here, we systematically functionalized three components of migration-permissive hydrogels to foster salivary tissue morphogenesis. We separately analyzed contributions of an enzymatically degradable crosslinker, a pendant integrin-binding site, and hydrogel porosity to best support high viability, integrin-dependent cell adhesion and migration. Structure size, frequency, and morphology were all affected by hydrogel crosslink density and integration of biofunctional peptides. Viability and proliferation data suggested that integration of integrin binding sites had the greatest effect on hS/PCs behavior. A larger internal matrix space, created by increasing both crosslinker length and PEG content, was needed to form large multicellular hS/PC structures. Peptide-modified hydrogels with more internal space shifted hS/PC organization from spheroidal, surrounded by thick basement membrane, to an asymmetric arrangement with punctate matrix proteins defining a “wrinkled” perimeter. Integrin-binding peptides activated integrin β_1_, with highest activation observed in hydrogels having both cleavable peptide and integrin ligand. The design parameters we prescribe allowed us to encapsulate hS/PCs in a humanized biomimetic hydrogel matrix able to support morphogenesis needed for salivary restoration.

## 1. Introduction

Head and neck cancer remains an important public health issue, with over 65,000 new cases reported in 2020 [1]. Most of those affected patients with locally advanced disease will receive a standard treatment including radiation therapy, or a combination of radiation therapy with chemotherapy [2,3]. Although advancements have been made in the precision of radiation therapy through methods like intensity-modulated radiotherapy (IMRT), there is still collateral damage of surrounding tissues, most notably the salivary glands [4–7]. In most cases, radiation-induced damage to the functional units of salivary gland tissue leads to a dramatic and permanent decrease in salivary flow that severely impacts the patient’s oral health and quality of life [8,9]. Unfortunately, only palliative care (artificial saliva substitutes, cholinergic sialogogues, etc.) is available for patients suffering from this feeling of perpetual “dry mouth,” or xerostomia, and even these solutions have undesired side effects [10–13]. Other promising non-pharmacological treatments, including gene therapy [14,15] and preservation/reintroduction of stem/progenitor cell populations, are in development but are not yet available for clinical use [16–18]. Yet, these same therapies face ongoing deliberation within the field, as appropriate stem cell signatures, and the persistence of a differentiated phenotype after radiation, remain active areas of debate and investigation [19].

Tissue engineering offers a promising solution, as this paradigm classically employs patient-derived cells (usually autologous, possibly allogeneic), organized within a support matrix to restore native function. Multiple laboratories, including our own, have contributed discoveries toward the broader goal of bioengineering salivary glands for replacement of irradiated, dysfunctional salivary glands. These contributions include integration of select cell populations and implementation in multiple biomaterial matrices [20–24]. Ultimately, clinical translation of these models will require a biocompatible matrix that will enable incorporation of the implanted neotissue. Although work with rodent-derived salivary cell lines has been instrumental for pathway discovery in salivary gland development, such discoveries must be reconfirmed with human cells for validation and translational use. Much remains to be uncovered regarding the mechanisms of salivary gland regeneration and the specific progenitor cell populations responsible for regenerating each compartment of this complex organ [25].

Our lab proposes to overcome the problem of hyposalivation by bioengineering an autologous, functional salivary gland from primary human salivary-derived stem/progenitor cells (hS/PCs). These have been reliably and successfully grown in our laboratories in long-term, 3D cultures within thiolated hyaluronic acid (HA)-based hydrogels, crosslinked with poly(ethylene glycol) diacrylate (PEGDA) [26]. Furthermore, we demonstrated that HA hydrogels, covalently modified with extracellular matrix (ECM)-derived, bioactive peptides, supported the preferential differentiation of hS/PCs into an acinar cell phenotype [27]. Although these soft, HA-based matrices permit hS/PC proliferation and organization into spheroids with discernible interior lumen, these structures do not progress further to the native organization of branched acini, wrapped by myoepithelial cells and basement membrane, coalescing in elongated ductal segments. HA-PEGDA matrices are not readily degradable by cells within physiologic timeframes and have restricted pore size within concentration ranges of typical use. These biocompatible matrices support large, viable, and relatively uniform spherical salivary structures, but not the asymmetric branched morphologies that are phenotypic of mature glands.

For the present work, we focused on customizing an HA-based scaffold with enzymatically-degradable crosslinks and integrin-binding ligands to enable cell motility and exploration of 3D space, both of which are needed for generation of an elongated, interconnected salivary structure *in vivo*. We individually varied each of three design parameters consisting of a cleavable crosslinker, integrin-binding site, and porosity. We hypothesized that the use of a degradable matrix with integrin ligands would encourage the organization of salivary-derived cells into more complex structures that ultimately would better resemble the early stages of SG formation, and advance our development of an engineered SG substitute.

## 2. Materials and Methods

### 2.1. Materials

Dulbecco’s Modified Eagle Medium (DMEM)/Nutrient Mixture F-12 (DMEM/F-12), penicillin/streptomycin solution, and Fungizone (amphotericin B), and trypsin-EDTA was purchased from Thermo Fisher Scientific, Grand Island, NY. Hepatocyte Culture Media Kit (item 355056) with supplemented epidermal growth factor (EGF) was purchased from Corning, Inc., Life Sciences, Oneonta, NY. Soybean trypsin inhibitor was purchased from Sigma-Aldrich, St. Louis, MO. Thiolated hyaluronic acid polymers (HA-SH) and PEGDA were purchased as a kit (HyStem ^®^) from ESI BIO, Alameda, CA. HA-SH was provided under as a lyophilized powder under inert gas blanket and slight vacuum by the supplier, with ~40% modification of carboxyl groups to thiol groups. All peptides used in this study were purchased from GenScript USA, Inc., Piscataway, NJ. N-(2-Hydroxyethyl)piperazine-N-(4-butanesulfonic acid) (HEPBS) was purchased from Santa Cruz Biotechnology, Inc., Dallas, TX. Acrylate-PEG-succinimidyl valerate (Ac-PEG-SVA) (3400 g/mol) was purchased from Laysan Bio, Inc., Arab, AL. Dialysis membrane (3500 Da MWCO) was purchased from Spectrum Laboratories, Inc., Rancho Dominguez, CA. Phosphate buffered saline (PBS) was purchased from Lonza, Walkersville, MD. Sylgard^™^ 184 poly(dimethylsiloxane) (PDMS) was purchased from Dow Corning, Midland, MI. Information for antibodies, counterstains and cell dyes are listed in supplementary information (Table S1 and Table S2).

### 2.2. Isolation of primary salivary-derived human stem/progenitor cells (hS/PCs) from parotid tissue

Salivary-derived human stem/progenitor cells (hS/PCs) were isolated from healthy margins of human salivary gland tissue of head and neck cancer patients undergoing tumor resection. Protocols were approved by institutional review boards at Rice University and University of Texas Health Science Center at Houston. Human parotid gland tissue was shipped immediately in sterile 50 mL conical tubes on ice for overnight delivery to our laboratories in Houston, TX. Tissue specimens were disinfected with betadine solution (1% vol/vol) in cold DMEM/F-12. Tissue was minced into a slurry and suspended in complete Hepatocyte culture medium supplemented with epidermal growth factor (EGF) (10 ng/mL), penicillin (100 U/mL)/streptomycin solution (100 μg/mL) and Fungizone (amphotericin B) (1% vol/vol) (“complete Hepatocyte culture medium”). Hepatocyte medium suspension with tissue pieces was plated into T-75 tissue culture flasks, enabling the tissue pieces to attach to the culture surface. Explants grew from each piece until cells reached 70%-80% confluency. Cells in each T-75 flask were passaged using 3 mL of trypsin-EDTA (0.05% wt/vol) for 5 min, or until cells detached from plates, followed by enzyme deactivation with 3 mL of soybean trypsin inhibitor (1 mg/mL). Recovered cells were resuspended in complete Hepatocyte culture medium, and plated into T-75 tissue culture flasks, where they were expanded. Experiments were repeated with cells isolated from three different female patients, ages ranging from 22-57. Primary hS/PCs passage 3 through 8 were used for these studies.

### 2.3. Acrylation of MMP-labile peptidic crosslinkers and pendant integrin ligands

The protocol for synthesis of acrylated peptides was reported previously [28,29]. Collagen I-derived, MMP-labile peptide KGGGPQG↓IWGQGK (“PQ”; ↓ denotes cleavage site) and MMP-insensitive peptide KGGG<u>D</u>QGI<u>A</u>G<u>F</u>GK (“PQC”) have been used previously for developing matrices supportive of cell migration, and their controls, respectively [30,31]. Both PQ and PQC peptides were synthesized commercially, acetylated at the N-terminus. Each peptide was reacted with Ac-PEG-SVA (3400 g/mol) at a molar ratio of 1:2.2 in HEPBS buffer (20 mM HEPBS, 100 mM NaCl, 2 mM CaCl_2_, and 2 mM MgCl_2_) at an adjusted pH of 8.0 using 0.1 N NaOH. Similarly, fibronectin-derived peptides G<u>RGD</u>S and scrambled G<u>RDG</u>S were synthesized commercially, with C-terminal amidation, then reacted with Ac-PEG-SVA (3400 g/mol) at a molar ratio of 1.2:1 in HEPBS buffer at a pH of 8.0. Reactions proceeded in an amber vial covered with foil for 12 hrs at 4°C under orbital shaking conditions. Acrylated peptides were transferred into a pre-washed 3500 Da MWCO dialysis membrane and dialyzed against 4 L of milliQ water under magnetic stirring conditions. MilliQ water was replaced five times over 2 days. Reacted peptide products were sterile filtered and lyophilized to a dry powder. Acrylated peptides were characterized by MALDI-TOF (Bruker AutoFlex II). Approximate molecular weights were: 8111 g/mol (Ac-PEG-PQ-PEG-Ac), 8033 g/mol (Ac-PEG-PQC-PEG-Ac), 3889 g/mol (Ac-PEG-GRGDS), and 3889 g/mol (Ac-PEG-GRDGS).

### 2.4. Preparation of molds

Methods for creating molds from poly(dimethylsiloxane) (PDMS) have been previously described in Fong et al. (2014) and Zarembinski et al. (2017) [32,33]. Briefly, large PDMS slabs were cast within metal plates at 1 mm thickness and cured in a Carver press at 150°C for 20 min at 1000 psi. Rectangular pieces of dimensions 24 x 60 mm were cut with a CO2 laser cutter (X-660, Universal Laser Systems, Scottsdale, AZ).Within each rectangular slab, multiple cylindrical cavities (6mm diameter) were cut as molds for hydrogels. PDMS molds and glass microscope slides were sterilized by steam-autoclaving. PDMS molds were laid onto glass microscope slides with sterile forceps and press-sealed to avoid leakage of the aqueous hydrogel solution. Individual pieces of mold-slide apparatus were placed in sterile petri dishes and were exposed to UV overnight.

### 2.5. Hydrogel preparation and hS/PC encapsulation

Hydrogel networks for base HA-PEGDA and peptide-modified PQC-RDG and PQ-RGD are illustrated in Figure 1. Working solutions of HA-SH were prepared in sterile degassed water at 10 mg/mL. Working solutions of pegylated pendant peptides (acrylate-PEG-G<u>RGD</u>S and acrylate-PEG-G<u>RDG</u>S) were prepared in PBS at 70 mg/mL or 18 mM. Working solutions of pegylated crosslinkers (Ac-PEG-PQ-PEG-Ac and Ac-PEG-PQC-PEG-Ac; varying concentrations listed in supplemental Table S3) were prepared in PBS. All hydrogels were formed using a volume ratio of 4:1:1 (working solutions of HA-SH: crosslinker: pendant peptide or PBS for HA-PEGDA hydrogels). The concentration of each crosslinker working solution was varied for various crosslinking densities to yield thiol:acrylate (SH:Ac) ratios of 6:1, 11:1, and 16:1, with corresponding final crosslinker concentrations of 0.46 mM, 0.25 mM, and 0.17 mM. The concentration of pendant peptides, if used, in each hydrogel was held constant, at a final concentration of 3 mM.

**Figure 1.**
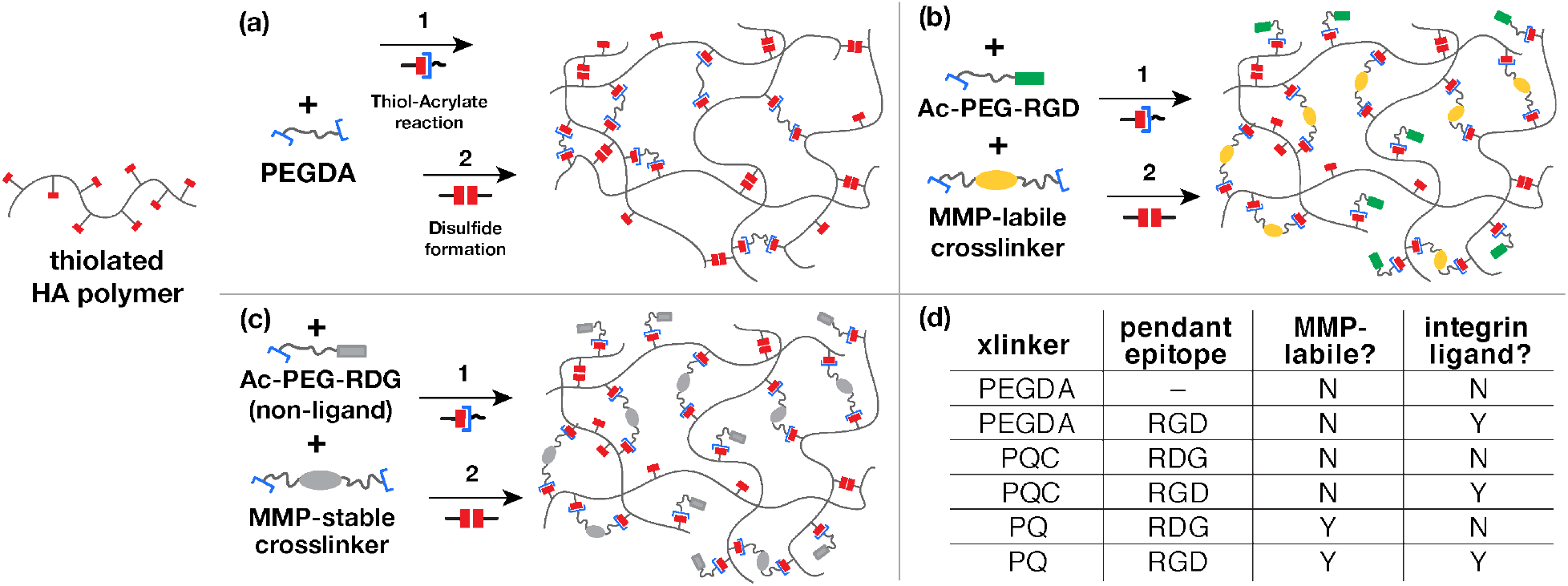
Schematic of HA-based hydrogels used for hS/PC culture. The base hydrogel matrix comprised thiolated HA and a series of PEG-based crosslinkers and pendant epitopes, covalently linked via a thiol-acrylate addition. (a) Low molecular weight PEGDA was used as a base reference hydrogel. This was modified by substituting PEGDA with either (b) a MMP-cleavable peptide crosslinker (“PQ”) and a pendant integrin-binding RGD site, or (c) an equimolar concentration of a control, MMP-stable peptide crosslinker (“PQC”) and a scrambled pendant RDG non-integrin-binding site. The table in (d) shows permutations of each hydrogel type, swapping crosslinkers or pendant epitopes, that are assessed in the paper.

For cell encapsulations, HA-SH working solution and acrylated pendant peptide solution (or PBS for HA-PEGDA) were first mixed and allowed to react for 30 minutes. Primary hS/PCs were trypsinized from tissue culture flasks as noted above, centrifuged into cell pellets at a defined cell number. Crosslinker working solutions were added to the above HA solutions, reacted for 5 minutes, and the total hydrogel solution was used to gently resuspend the prepared cell pellets until the hydrogel suspension was uniform. Cell density in the final hydrogel solution was 2.4E6 cells/mL. Hydrogel-cell suspensions (42 μL/ea) were plated into cylindrical cavities in the molds and allowed to crosslink for 30 min at 37°C in a humidified incubator. A 40μL drop of complete Hepatocyte culture medium was then gently added onto each hydrogel. Hydrogels were allowed to further crosslink at 37°C for an hour, then a 25G syringe needle was used to trace the periphery of the hydrogels to release them from the wall of the PDMS molds. PDMS molds were carefully removed from glass slide using sterile tweezers. Hydrogels were detached from the glass microscope slides using stainless steel spatulas and were placed into 48-well plates containing 500 μL of complete Hepatocyte culture medium. Plates containing hydrogels were placed into 37°C incubators for the length of individual experiments. All hydrogels were replenished with 300 μL of fresh medium every three days.

### 2.6. Measurement of hydrogel size and swelling

Cell-laden hydrogels were removed from medium at day 1 after encapsulation and placed in the center of wells in a 6-well plate. Excessive medium surrounding hydrogels was removed with a P200 micropipette. Hydrogels were measured with a caliper at day 1 after encapsulation. Means were calculated from n=3 hydrogels.

### 2.7. Viability assay on encapsulated hS/PCs

Viability assay was performed at days 1, 7, 14, and 25. Viability assay solution consisted of 2 μM calcein AM, 4 μM ethidium homodimer III (EthD-III), and 10 μg/mL of Hoescht 33342 (listed in supplemental Table S2) in complete Hepatocyte culture medium. Hydrogels were placed in a new 48-well plate, and each hydrogel was treated with 300 μL of viability assay solution at 37°C for 30 min, and then imaged using a Nikon A1-Rsi confocal microscope (Plan Apo λ 20x, 512 x 512, 1.375 μm step size, 150 μm Z-stack depth). Viability assays were repeated on encapsulated cells from three different female patients; samples from two female patients were quantified. A minimum of n=3 hydrogels per group with triplicate Z-stacks per hydrogel was used. Single frames were processed with NIS-Elements AR software (Nikon Instruments, Melville, NY) and Z-stacks were quantified via Imaris software version 9.2.1 (Bitplane USA, Concord, MA). Quantification details are provided in methods section 2.9.

### 2.8. Immunofluorescence and confocal imaging of hS/PCs

Hydrogels were harvested at various time points, fixed with 4% paraformaldehyde in PBS for 15 min at room temperature, and stored in PBS at 4°C prior to immunostaining. Hydrogels were cut into fourths, and each segment was placed into individual wells in 96-well plate. Cells were permeabilized by immersion in 0.2% Triton X-100 in PBS for a 15 min incubation period. Blocking of hydrogels was performed with filtered 3% goat serum (GS) in PBS solution overnight at 4°C. All primary and secondary antibody staining was conducted overnight at 4°C, in filtered 1% GS/0.2% Triton X-100 in PBS, and all are listed in supplemental Table S1. Primary antibodies were used at a 1:100 dilution, and secondary antibodies were used at a 1:500 dilution, from the manufacturer, with PBS rinsing between steps. Cells were counterstained with DAPI and fluorophore-tagged phalloidin in PBS for 30 min at room temperature; concentration details are listed in supplemental Table S2. A minimum of n=3 Z-stacks per group were used. Samples were imaged using a Nikon A1-Rsi confocal microscope (Apo LWD 40x WI λS DIC N2, 1024 x 1024, 0.675 μm step size, Z-stack depth sizes used for quantification were the same for all groups). Single-frame images were created with NIS-Elements AR software (Nikon Instruments, Melville, NY) and Z-stacks were quantified via Imaris software version 9.2.1 (Bitplane USA, Concord, MA). Quantification details are provided in methods section 2.9.

### 2.9. Image analysis of Z-stacks

All Z-stacks were analyzed and quantified in Bitplane Imaris software. Cell viability assays were quantified using the Imaris Spots function to identify and count all Hoechst^+^ nuclei with a maximum diameter of 8 μm and all nonviable EthD-III ^+^ cells with a maximum diameter of 12 μm. Cell viability percentages were calculated by following equation:

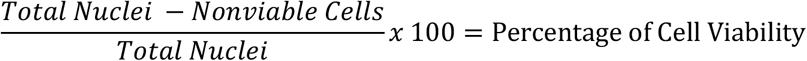

Mean cell viability percentage for n=3 Z-stacks per condition (estimated 300-600 total cells per stack) with standard deviation is shown in column graphs.

Volumes and sphericity for multicellular structures were quantified from viability assays using the Imaris Surfaces function. Surfaces were identified around calcein^+^ structures, encompassing three or more Hoechst^+^ nuclei, and the volume contained within each surface was calculated. Sphericity was calculated by the Imaris software using the following equation:

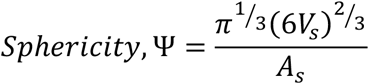

where *V_s_*= structure volume, and *A_s_* = structure surface area. The total number of multicellular structures formed in each Z-stack at day 7, day 14 and day 25 were quantified using the same methods for quantifying structure volumes.

Proliferation was quantified by counting the number of Ki67^+^ nuclei in Z-stacks from Ki67 immunostaining. An object was created using the Surfaces function for each nucleus with Ki67 expression above a manual intensity threshold of 500 a.u. Total nuclei were quantified the same way that they were quantified for cell viability percentages; however, DAPI counterstain was used instead of Hoechst. Percentage of Ki67+ cells per group for n=3 Z-stacks was then graphed into columns with mean and standard deviation.

Activated integrin β_1_ expression per cell was quantified using Z-stacks from immunostaining. The Surfaces function was used to create a region of interest encompassing all integrin β_1_ expression around each cell above an intensity threshold of 200 a.u. Values for average intensity sum per cell for each sample (n=3 hydrogels with triplicate Z-stacks) were then graphed into columns showing mean and standard deviation.

### 2.10. Rheological Characterization of Hydrogels

Rheological characterization of cell-free hydrogels was performed using a TA Instruments DHR-3 rheometer with a 20mm stainless steel parallel plate geometry, at a 1mm gap distance, on a temperature-controlled Peltier Plate base. A water-filled solvent trap and evaporation cover were used to reduce evaporation. Aqueous components of each hydrogel variant were prepared individually as described above for cell encapsulation, mixed, and immediately loaded within the plate geometry. A thin layer of mineral oil was applied carefully around the exterior of the hydrogel to further reduce evaporation. An initial strain sweep, from 10^-2^ to 10^2^ oscillation strain (%), was conducted to confirm that all experiments would be conducted within a region of linear viscoelastic response. Time studies were performed with 25% strain, 1 Hz frequency, at 37°C for 6hrs. Two separate specimens were measured for each hydrogel variant of composition and crosslink density.

### 2.11. Hydrogel Degradation with Collagenase Type I

Three hydrogel variants, crosslinked at 11:1 (SH:Ac), were prepared in triplicate within PDMS molds as described above, then incubated in 1XPBS at 37°C for two days to equilibrate. Each was then incubated in collagenase type I solution (1μg/mL) in 1XPBS at 37°C for six days. Hydrogel diameters were measured daily with calipers over the duration of the experiment, and reported as an average fold change in diameter.

### 2.12. Statistical Analysis

Statistical analysis for hydrogel measurements and structure sphericity was performed via a one-way ANOVA with Dunnett’s post-hoc test between each group and HA-PEGDA. Statistical analysis for hydrogel degradation measurements was performed via a two-way ANOVA. Statistical analyses for cell viability, multicellular structure sizes, percentage of proliferating cells, and activated β_1_ integrin expression were performed by a one-way ANOVA with Tukey post-hoc test between each group. Statistical analysis for the number of multicellular structures formed was performed via a student’s t-test between hydrogels crosslinked with the same crosslinker with or without RGD at each timepoint. A p-value <0.05 was considered significant. p-values: * = p<0.05; ** = p<0.01; *** = p<0.001; **** = p<0.0001.

## 3. Results

### 3.1. Growth and viability of hS/PCs in customized HA matrices

To assess the impact of migration-permissive hydrogels on hS/PC growth and viability, hS/PCs were encapsulated as single cells in base HA hydrogels (HA-PEGDA), and in multiple formulational permutations of these gels, through covalent network inclusion of the integrin ligand RGD (or a scrambled control), and an MMP-labile peptide-based crosslinker (or its MMP-stable analog). These variants are illustrated in Figure 1. The susceptibility of hydrogels containing the “PQ” crosslinker to enzymatic degradation was confirmed through exposure to collagenase type I (Figure S1). Migration-permissive PQ-RGD hydrogels gradually expanded in size with continued enzyme exposure, while PQC-RDG and HA-PEGDA gels were unaffected. To further assess effects on cell growth and viability associated with changes in the surrounding ECM network properties (swelling, stiffness and porosity) perceived by cells, we varied the crosslinking densities (thiol to acrylate ratios; SH:Ac) of each hydrogel type by adjusting the molar concentration of crosslinkers, while keeping a constant molar concentration of pendant peptides in peptide-modified hydrogels. Notably, the identified SH:Ac ratios (6:1, 11:1, and 16:1) refer only to molar concentration of acrylates from crosslinkers and do not include the additional acrylate contributions of pendant peptides.

**Supplementary Figure S1.**
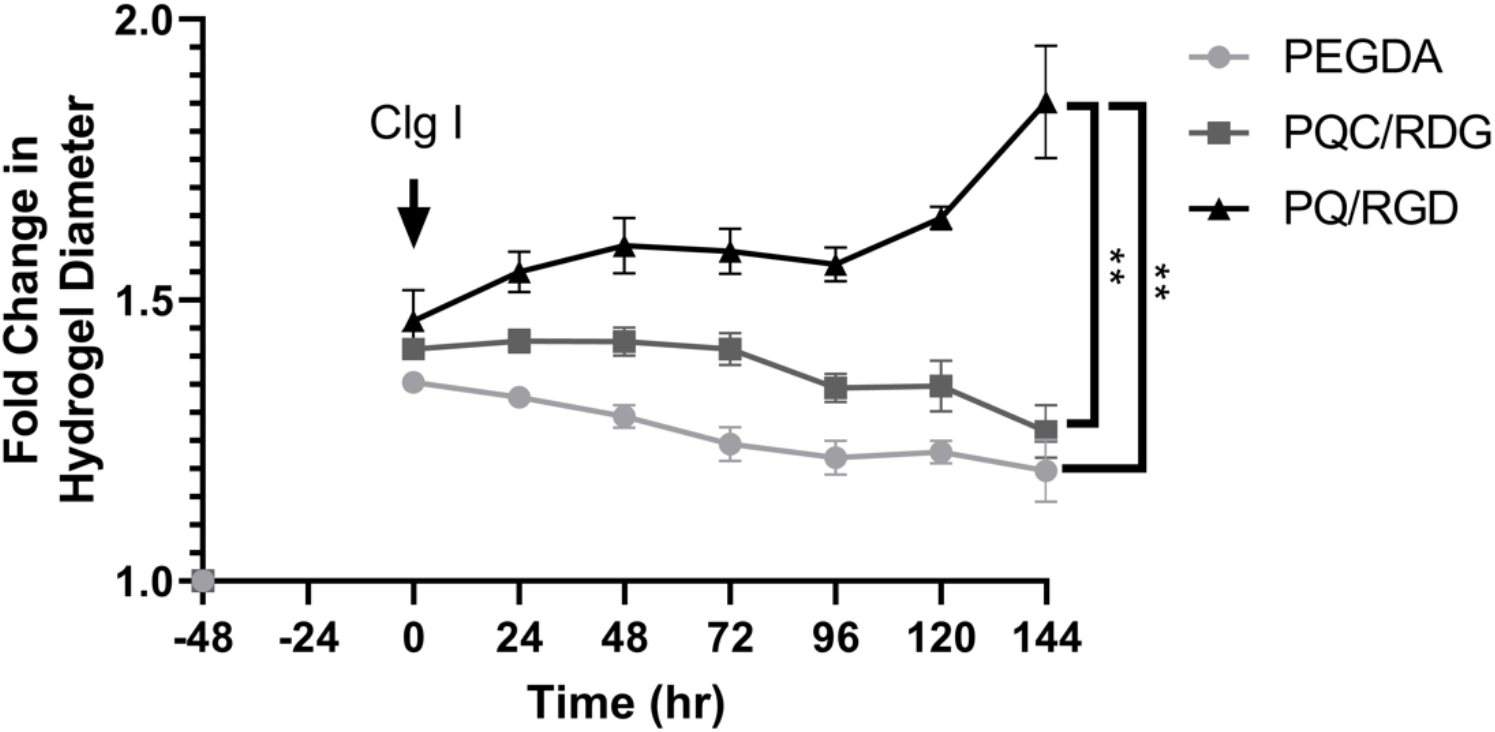
Hydrogel response to collagenase incubation. Listed hydrogels (11:1; SH:Ac) were prepared by standard methods, released from molds, and incubated in 1XPBS at 37°C for 48 hrs. At t=0, hydrogels (n=3 per variant) were incubated in a solution of collagenase type I (1μg/mL) in 1XPBS at 37°C and the diameters of each were measured daily for six days. Average fold change in diameter of each hydrogel type over time is shown; error bars indicate standard deviation. Statistical analysis was performed via a two-way ANOVA.

The viscoelastic properties and gelation profiles of the HA-PEGDA, PQ-RGD, and PQC-RDG HA-based hydrogels at the various crosslinking densities were characterized (Figure S2). After loading onto the rheometer, all formulations quickly showed a G’-G” crossover indicative of hydrogel formation via crosslinking. Each reached a plateau storage modulus (G’) value within 10 min, and the final G’ value varied with crosslink density. As expected, PQ-RGD and PQC-RDG hydrogels at equivalent crosslink densities had comparable G’ values. Peptide-modified hydrogels had a consistently higher G’ value than their HA-PEGDA counterparts, at an equivalent crosslink density.

**Supplementary Figure S2.**
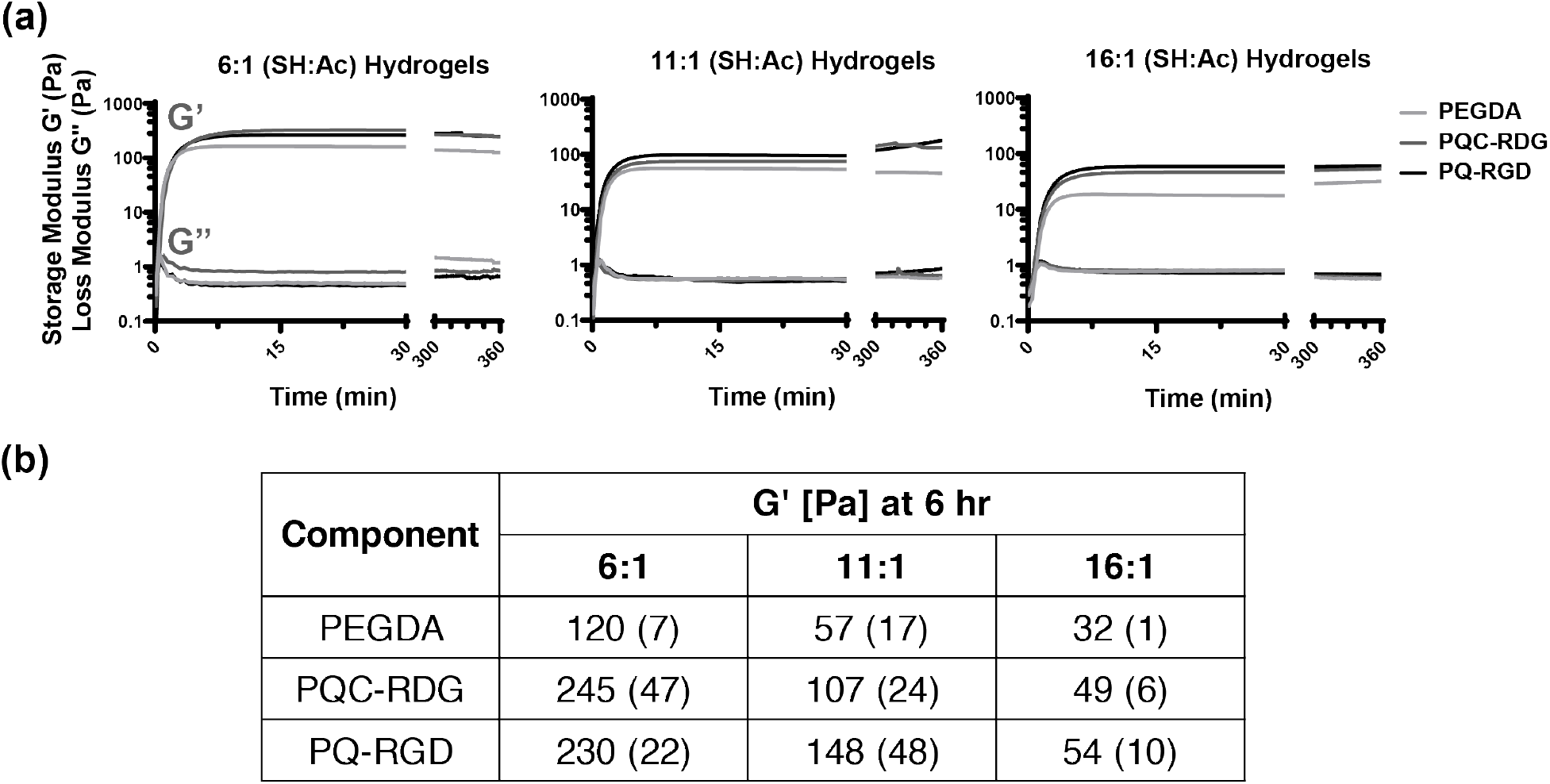
Rheological characterization of hydrogels over time. Time sweeps were performed on hydrogels immediately after mixing components to measure storage (G’) and loss modulus (G”) over a 6hr time course. (a) The three types of hydrogels crosslinked at 6:1, 11:1, and 16:1 (SH:Ac) were tested and compared to each other within each crosslinking density. Representative curves are shown from n=2-3 experiments. (b) Values for average (standard deviation in parentheses) of G’ at the final timepoint of 6 hr are listed.

Viability and growth of hS/PCs encapsulated within the multiple hydrogel variants were assayed at multiple timepoints (D1, 7, 25) (Figures 2, S3). High viability generally was maintained in all HA-based hydrogels at various crosslinking densities, except for a slight reduction in the base HA-PEGDA hydrogel at the highest crosslinking density (6:1) (Figure 2a). Highly viable (≥90%) multicellular structures were formed in all matrices by day 7 (Figure S3) and viability was maintained through day 25 (Figure 2b). Beyond this slight difference in viability at early timepoints, it was observed that the hydrogel variants induced a substantial difference in the size of multicellular hS/PC aggregates over time. In single frame images from viability assays performed at day 25, the largest multicellular structures found in peptide-modified hydrogels had a diameter of ~200μm, as compared to those in HA-PEGDA hydrogels, with a maximum diameter of ~90μm. Confocal Z-stacks of the viable multicellular clusters within each hydrogel were reconstructed within Imaris software, and individual calcein^+^ clusters were individually identified using the Surfaces function, generating the volume data shown in Figure 3a. Notably, the more conventional measure of structure diameter is not reported, because the structures in peptide-modified hydrogels were not only larger than those in base HA-PEGDA hydrogels, but also adopted a far less spherical morphology.

**Figure 2.**
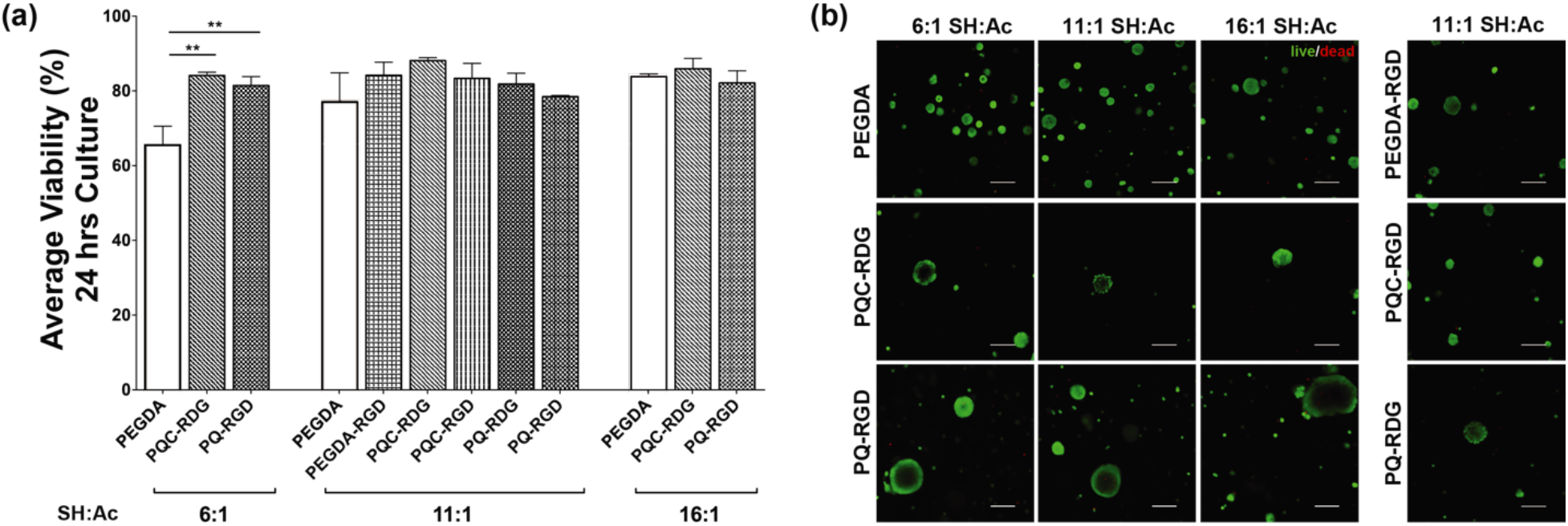
Viability imaging for hS/PC clusters in hydrogel variants. Viability assays were performed on hS/PCs encapsulated in HA-PEGDA, or the peptide-modified variants described in Figure 1d, across multiple timepoints of culture. Crosslink density in select gels was varied by altering the SH:Ac ratio, as shown. (a) HA hydrogels generally preserved high cell viability across three SH:Ac ratios after 24 hrs of culture, with no difference due to composition, except at the highest crosslink density (6:1). (b) Representative single-frame images are presented, out of n=4 hydrogels, after 25 days of culture. Peptide-modified hydrogels enable hS/PCs to grow into larger viable structures with lumen-like cavities. Viable cells are green, non-viable cells are red. Statistical analysis for (a) used one-way ANOVA with Tukey post-hoc test between each group. Scale bars = 100μm.

**Figure 3.**
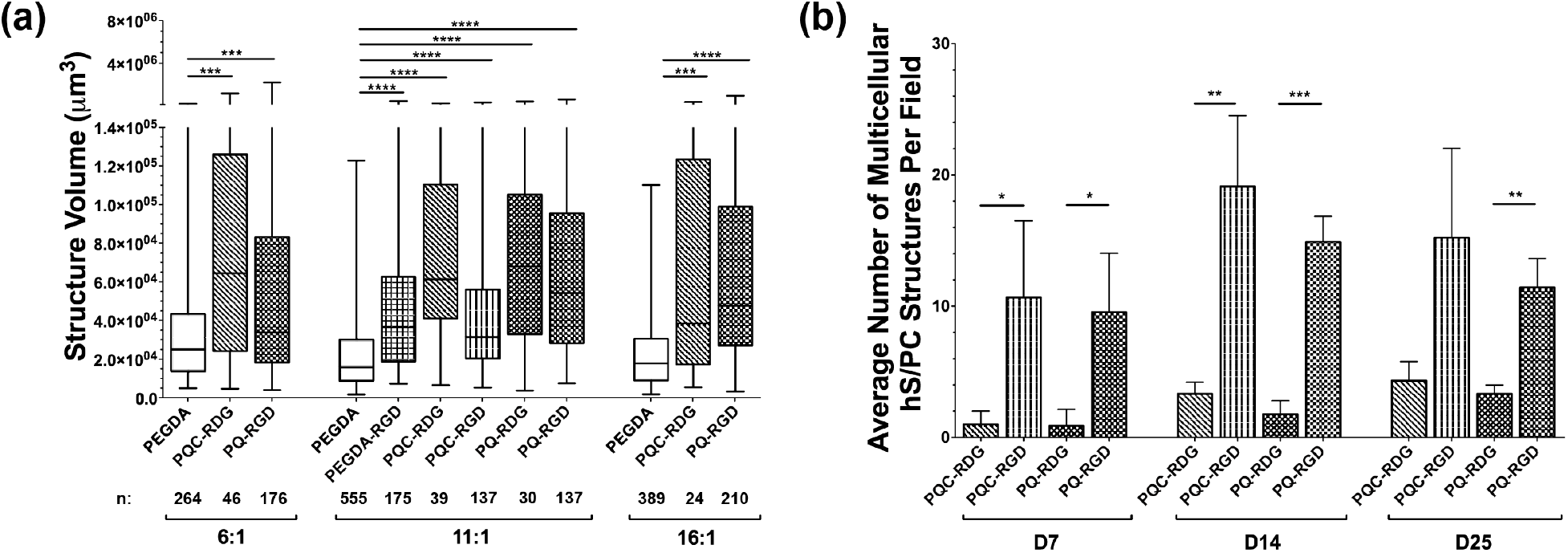
Characterization of hS/PC cultures in HA-based hydrogels. (a) Peptide-modified hydrogels supported larger multicellular structures than base HA-PEGDA networks. Number of clusters assessed per variant is reported under graph. (b) Integration of an RGD integrin-binding site led to a significant increase in the number of structures formed over time. n ≥ 3 hydrogels for all conditions. Statistical analysis for (a) used one-way ANOVA with Tukey post-hoc test among each group; (b) used a student’s t-test between compositions with the same crosslinker, with or without RGD. (a) Graph shows a box/whisker plot with median, 1^st^/3^rd^ quartiles, and max/min. Error bars in (b) are SD.

**Supplementary Figure S3.**
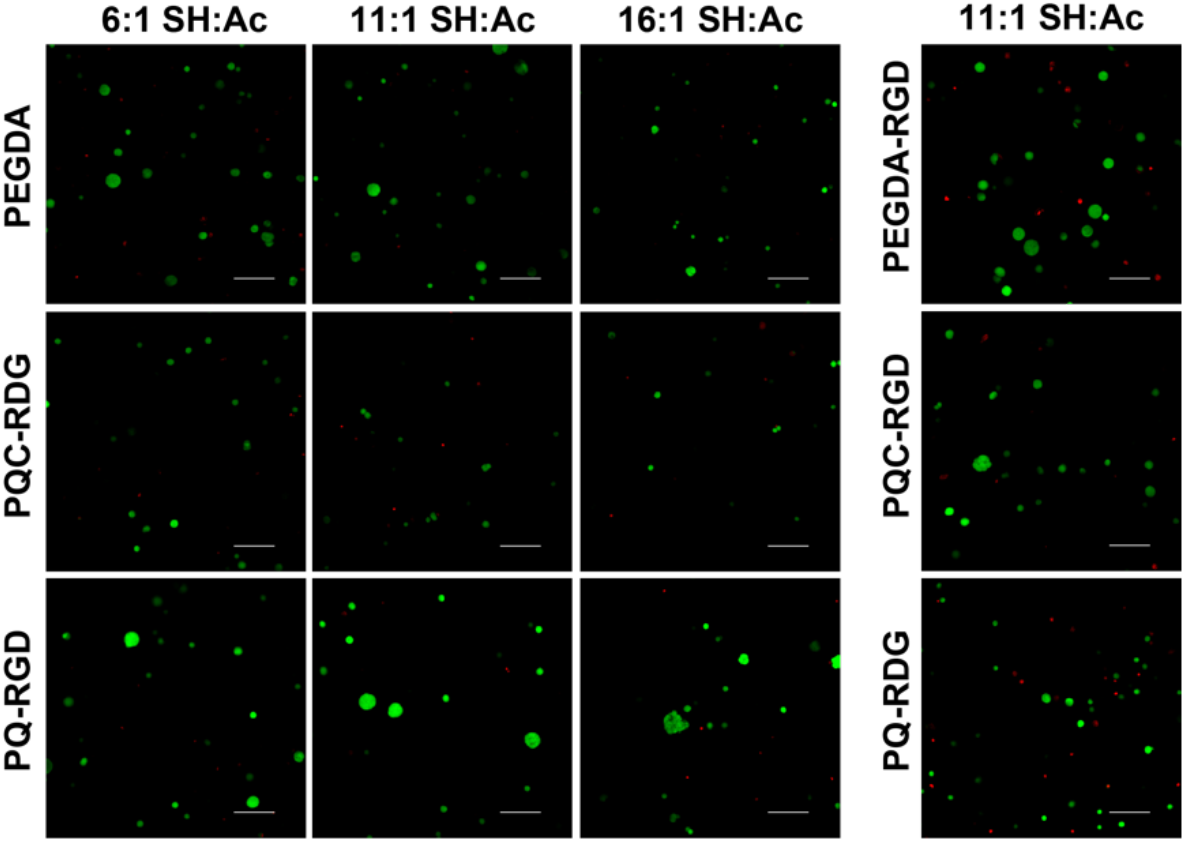
Viability imaging for hS/PCs in hydrogel variants. Viability assays were performed on hS/PCs encapsulated in HA-PEGDA, or the peptide-modified variants described in Figure 1d, after 7 days of culture. Crosslink density in select gels was varied by altering the SH:Ac ratio, as shown. Representative single-frame images are presented, out of n=4 hydrogels. Peptide-modified hydrogels containing integrin-binding sites promoted multicellular structure formation by day 7. Viable cells are green, non-viable cells are red. Scale bars = 100μm.

The average number of multicellular structures per image field was quantified, as shown in Figure 3b. (Notably, the base PEGDA-crosslinked hydrogels were excluded from this comparison, as those matrices were substantially smaller after crosslinking, resulting in more visible structures per Z-stack, despite an equivalent starting cell density (Figure S4).) For the peptide-modified hydrogels, those with the pendant integrin-binding site RGD, and crosslinked with either noncleavable (PQC) or cleavable (PQ) (11:1; SH:Ac) crosslinkers, led to a significantly larger number of structures when compared to RDG hydrogels starting by day 7 and continuing through the last timepoint, day 25 (Figure 3b). Only the choice of integrin-binding site, and not the enzymatically-degradable crosslinker, led to increased frequency of multicellular hS/PC structures. Parallel studies were performed with separately-derived hS/PCs from three donors, and yielded qualitatively similar results for multicellular structure size, number, and morphology (Figure S5).

**Supplementary Figure S4.**
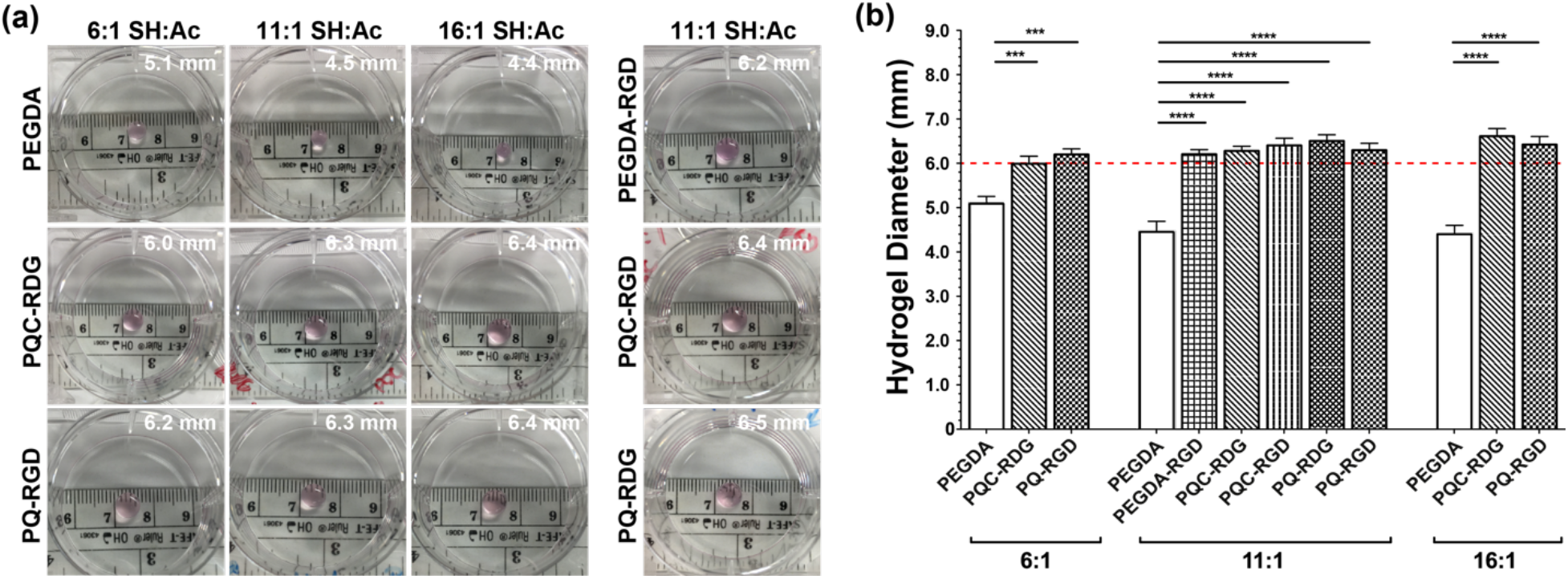
Hydrogel swelling ratios. (a) Representative images of cell-laden hydrogel variants, and measured diameters 24 hours after standard preparation. (b) Quantification of measurements, presenting averages of triplicate samples; error bars indicate standard deviation. Red dashed line indicates the diameter of the mold. Peptide-modified hydrogels swelled beyond the original mold size, while HA-PEGDA base hydrogels reduced in diameter after release from the PDMS mold. Statistical analysis was performed via a one-way ANOVA with a Dunnett’s post-hoc test between each group and HA/PEGDA.

**Supplementary Figure S5.**
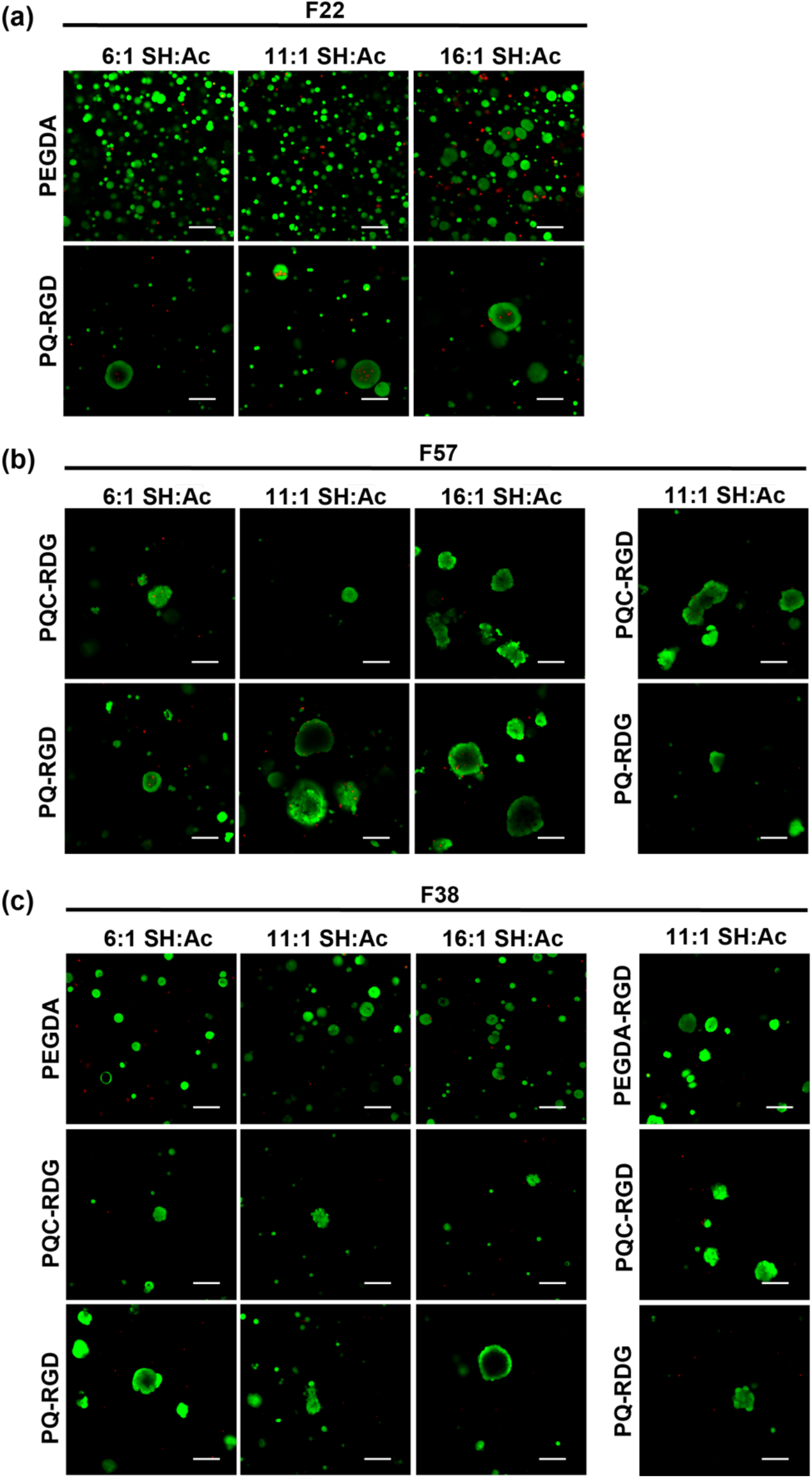
hS/PC response in hydrogels, across multiple patients. hS/PCs were isolated from three patients, all female, ages 22 (F22), 38 (F38), and 57 (F57). hS/PCs were encapsulated in HA-based hydrogels, cultured for 25 days, and assessed for qualitative viability and morphology. Representative images from live/dead staining show that hS/PCs responded similarly across patients. Scale bars = 100μm.

hS/PC structures in the various hydrogels were immunostained for proliferation marker Ki67 to assess hydrogel impacts on proliferation (Figure 4). Matrices containing an RGD integrin-binding site supported a significantly higher number of proliferating cells at day 14 compared to the control compositions without integrin-binding sites (Figure 4b). Proliferation of hS/PCs was unaffected when PQ was substituted for PQC in hydrogels lacking RGD.

**Figure 4.**
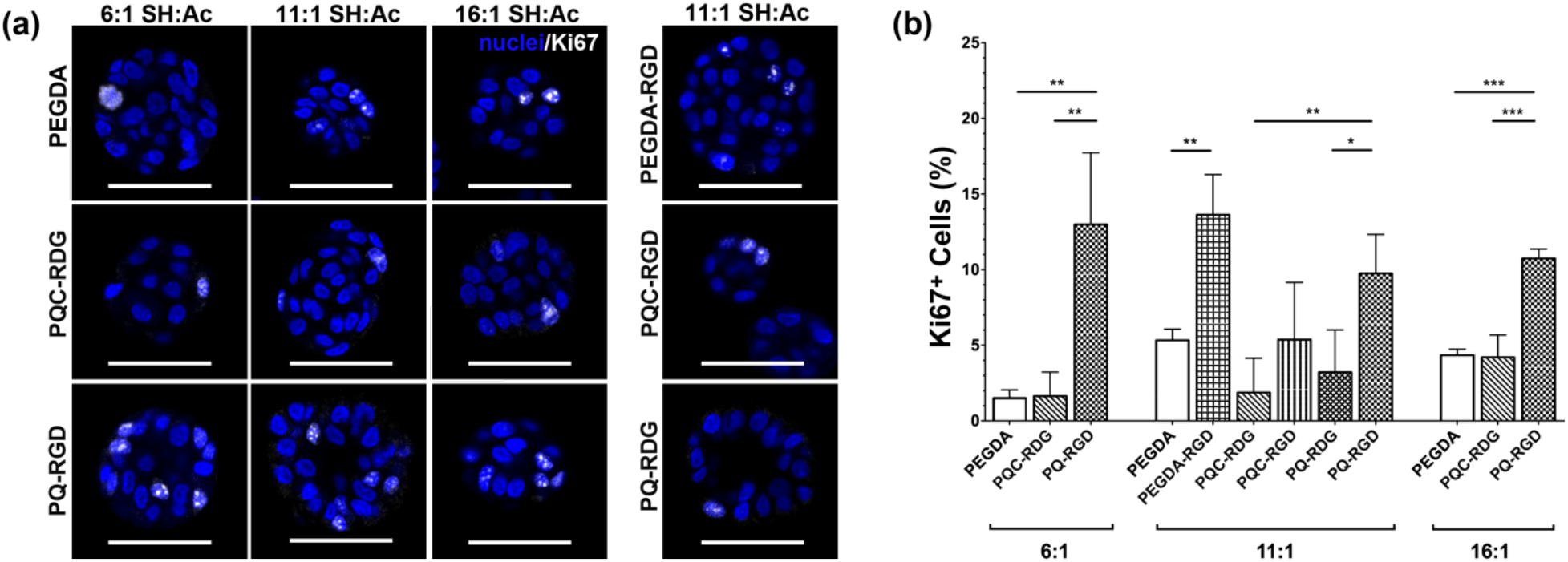
hS/PC proliferation in hydrogels. (a) Representative single-frame images show Ki67 immunostaining (white) of hS/PC multicellular structures within hydrogel variants. (b) Quantification of full Z-stacks identified that hydrogels containing integrin-binding site RGD consistently supported the highest levels of cell proliferation. n of at least 3 Z-stacks for each group. Nuclei are stained blue. Statistical analysis was performed using one-way ANOVA with Tukey post-hoc test between each group. Error bars are SD. Scale bars = 50μm.

### 3.2. Assessment of hS/PC interaction with surrounding matrix

hS/PCs in all three matrices were immunostained with an antibody that recognizes a ligand induced binding site (LIBS) of integrin β_1_ (activated), with enhanced recognition of the LIBS in the presence of fibronectin ligands (fragments or RGD) [34], to assess interactions between the cells and their surrounding matrix (Figure 5a). Because these cells begin synthesizing their own basement membrane soon after encapsulation (Figure S6), an early timepoint (day 1) was used to preferentially identify integrin activation due to the covalently-coupled ligands in the surrounding hydrogel. The volume and intensity of activated integrin β_1_ expression was integrated for hS/PCs in all matrices (Figure 5b). hS/PCs in PQ-RGD had the greatest activation of integrin β_1_ compared to the control variants at all three crosslinking densities. When we assessed the effect of incorporating each biofunctional peptide on integrin β_1_ activation (11:1; SH:Ac), it was observed across all matrices that integrating RGD always led to higher levels of activated integrin β_1_ than for equivalent hydrogels containing either a control RDG ligand, or no pendant ligand (Figure 5b). Activated integrin β_1_ levels were greater in hydrogels crosslinked with the longer crosslinkers (PQC or PQ), and thus more internal 3D space, compared to hydrogels crosslinked with the smaller crosslinker (PEGDA), and thus less internal 3D space, in the presence or absence of RGD. However, only the increase in integrin β_1_ activation associated with RGD integration instead of RDG in hydrogels crosslinked with PQ, and not those crosslinked with either PEGDA or PQC, was significant (p <0.05). Consequently, although RGD integration alone led to an increase in activated integrin β_1_, concurrent use of PQ and RGD led to the greatest increase in activated integrin β_1_.

**Figure 5.**
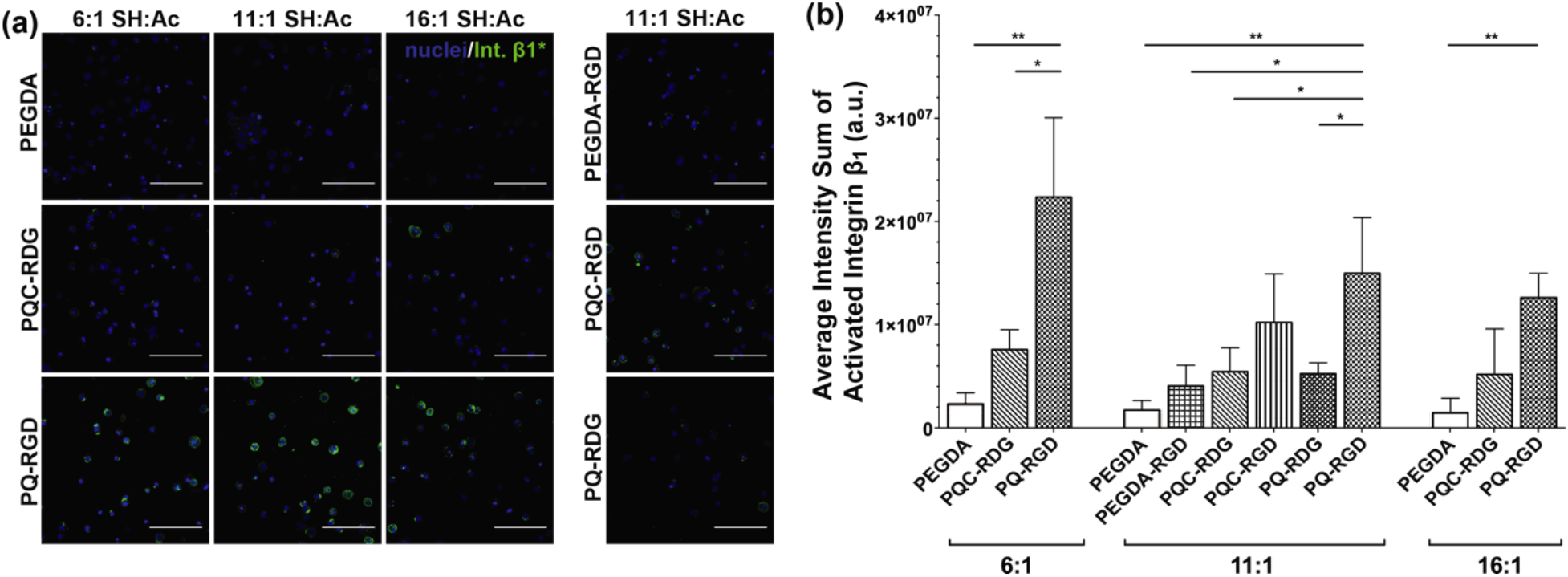
Levels of activated integrin β_1_ in hS/PC cultures. (a) Representative single-frame images show immunostaining for activated integrin β_1_ (green) around hS/PC multicellular structures within hydrogel variants. (b) Quantification of full Z-stacks identified that activation of expressed integrin β_1_ was highest in migration-permissive (PQ-RGD) hydrogels, compared to all other hydrogel variants. n=3 hydrogels for each group. Statistical analyses were performed using one-way ANOVA with Tukey post-hoc test between each group. Error bars are SD. Scale bars = 100μm.

**Supplementary Figure S6.**
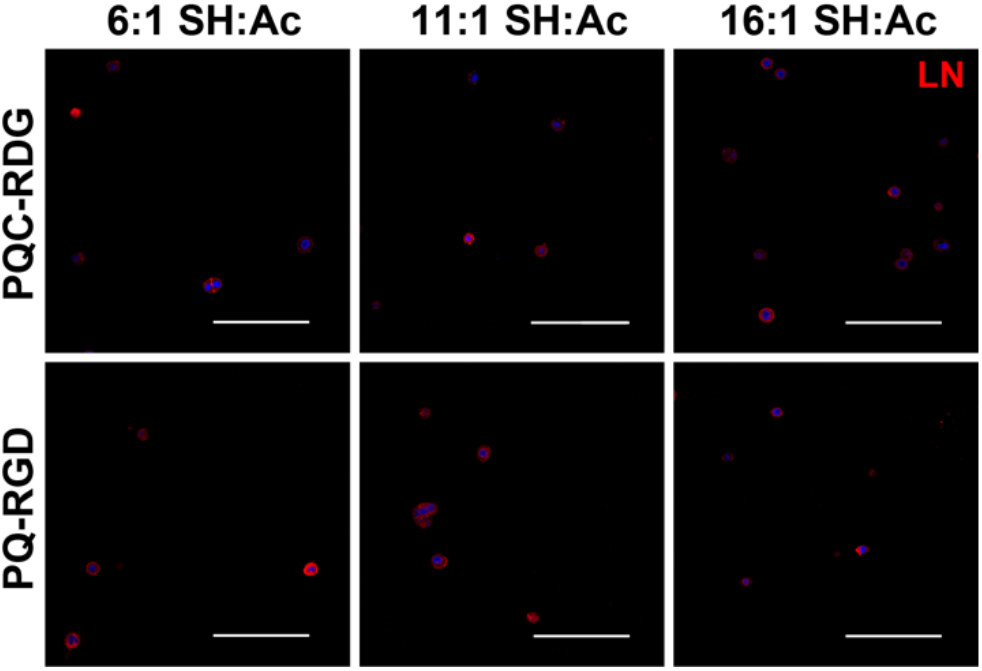
Early laminin expression by hS/PCs. Representative single-frame images show immunostaining of hS/PCs encapsulated in peptide-modified hydrogels for laminin (LN) at day 1. Single hS/PCs express LN at an early timepoint in all of hydrogel variants observed. Scale bars = 100μm.

To further assess hS/PC interaction with surrounding matrix, encapsulated hS/PCs at day 1 were immunostained for fibronectin receptor integrin α_5_ and laminin receptors integrins α_1_ and β_4_ (Figure S7a). Levels of each were highest in migration permissive hydrogels. Conversely, for hS/PCs cultured for the same period of time in migration impermissive hydrogels, laminin receptor proteins were not detected, and fibronectin receptor integrin α_5_ was identified but at very low levels. These observations support the data in Figure 5, and identify the influence of the matrix ligands at early timepoints.

**Supplementary Figure S7.**
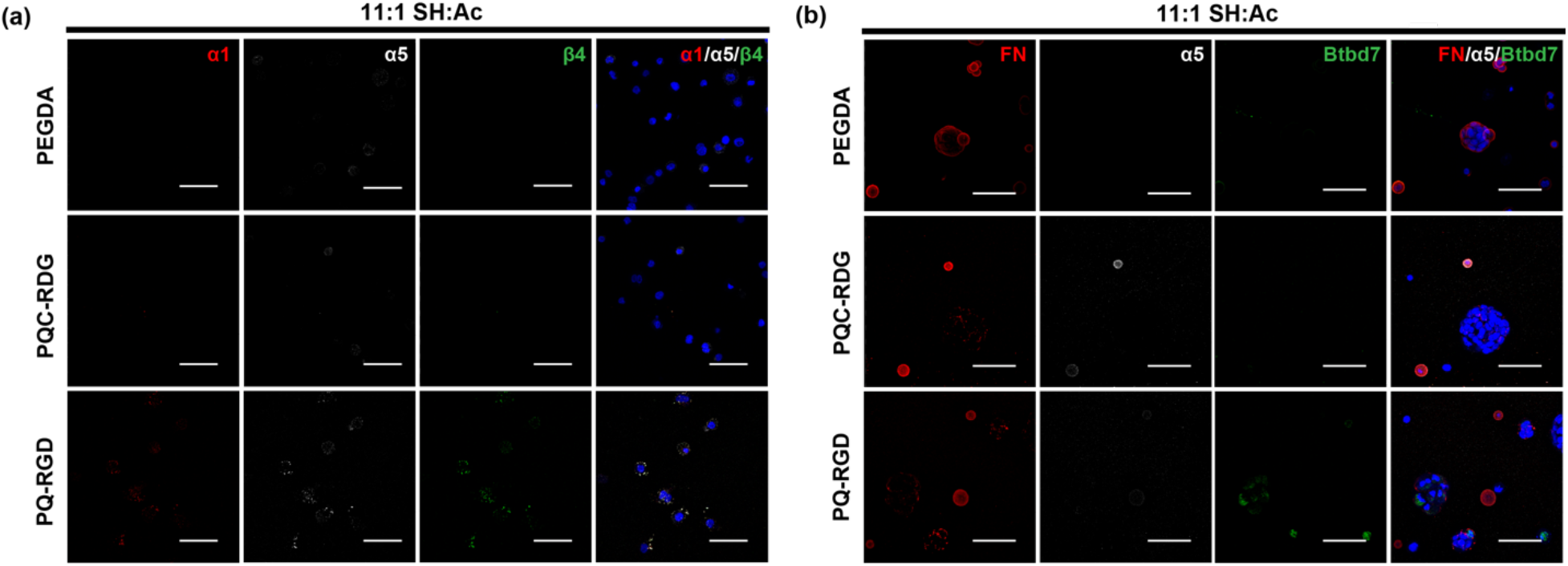
hS/PC interaction with extracellular matrix associated with clefting. (a) hS/PCs in three hydrogels (11:1 SH:Ac) were immunolabeled for laminin (LN) receptors (α1 [red] and β4 [green] integrins) and fibronectin (FN) receptor (α5 [white] integrin) at day 1 after encapsulation. LN receptors α1 and β4 integrin were almost exclusively expressed in PQ-RGD hydrogels. FN receptor α5 integrin was identified in all three hydrogel types, but levels were highest in PQ-RGD hydrogels. (b) To assess for impacts of the matrix after longer culture, hS/PCs in the hydrogel variants were immunostained for FN (red), FN receptor α5 (white) integrin and Btbd7 (green) at day 14. As was observed in Figure 7, FN was deposited uniformly as a thick band around the hS/PC spheroids in PEGDA hydrogels, and was punctate surrounding the “wrinkled” hS/PC clusters in the peptide-modified hydrogels. α5 integrin levels were low in the peptide-modified hydrogels, and absent in PEGDA hydrogels. Btbd7 expression by hS/PCs was observed in PQ-RGD hydrogels, particularly at the periphery of the multicellular clusters. No Btbd7 expression was detected in PEGDA or PQC-RDG hydrogels. (a-b) Images are maximum projections of individual Z-stacks, representative of staining from n=3 hydrogels. Scale bars = 50μm.

The morphologies of hS/PC structures in all hydrogel variants were compared to assess the impacts of matrix parameters on cell organization. hS/PC structures in HA-PEGDA self-organized into smooth spheroids, as has been observed in previous work, while structures in peptide-modified hydrogels were shaped more irregularly with cleft-like invaginations (Figure 6a). “Sphericity” is one tool that can assess for the degree of these invaginations, by ratioing contributions from structure volume to surface area [35]. Quantification of this parameter demonstrated that there was a shift of structure morphology from highly spherical (~1.0) in HA-PEGDA to significantly less spherical (<1) in peptide-modified hydrogels (Figure 6b). Representative volume reconstructions of the structures imaged in each Z-stack further demonstrated that migration permissive PQ-RGD hydrogels better supported the formation of asymmetric structures with lumen-like cavities and multiple cleft points than did HA-PEGDA matrices (Figure 6c).

**Figure 6.**
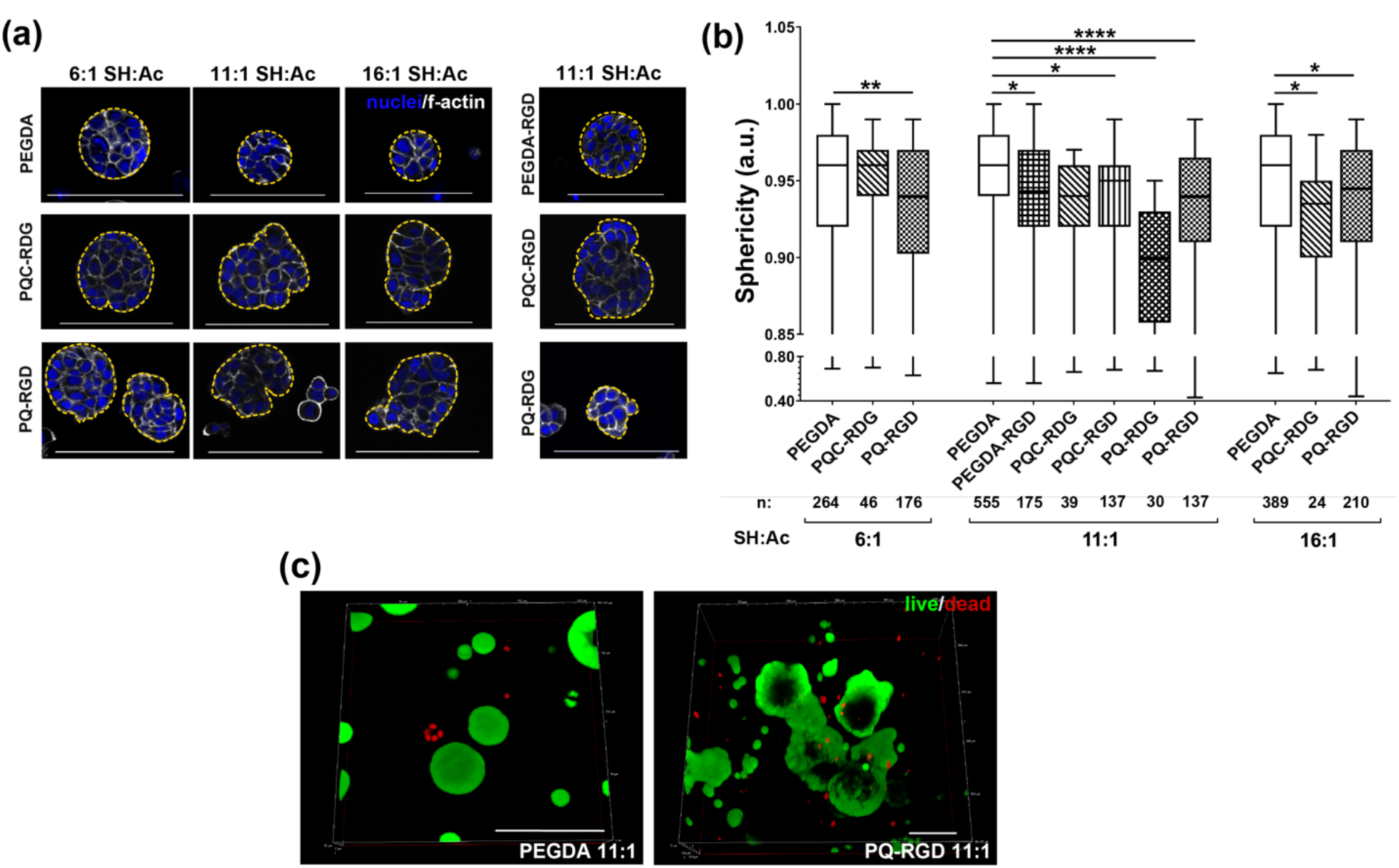
Multicellular structure morphology in hydrogel variants. (a) Representative single-frame images show fluorescent staining for F-actin (white; nuclei blue) in hS/PC multicellular structures within hydrogel variants at day 14; the periphery of each structure is highlighted in yellow. hS/PCs in HA-PEGDA gels were consistently spheroid, while those in peptide-modified hydrogels adopted a “wrinkled” periphery. (b) Sphericity values, measured for every cluster within each Z-stack, was highest in PEGDA gels, and significantly lower in most peptide-modified variants. Graph shows a box/whisker plot with median, 1^st^/3^rd^ quartiles, and max/min. Number of clusters assessed per variant is reported under graph. Statistical analyses were performed using one-way ANOVA with Dunnett’s post-hoc test between each group and HA-PEGDA. (c) Reconstructed 3D images from viability assays at day 25 demonstrate largely spheroidal structures within PEGDA hydrogels, while PQ-RGD variants enable more complex, asymmetric morphologies. n ≥ 3 hydrogels for all conditions. Scale bars in (a) = 100μm.

### 3.3. Basement membrane expression by hS/PC structures

Multicellular structures in base HA-PEGDA and peptide-modified hydrogels (11:1, SH:Ac) were immunostained for ECM proteins laminin and fibronectin to identify any changes in their expression pattern at day 14 (Figure 7). In HA-PEGDA hydrogels, a thick, even band of laminin was observed surrounding each spherical hS/PC structures (Figure 7a). Structures in peptide-modified, migration-impermissive (PQC-RDG) hydrogels maintained laminin expression hotspots dispersed inside and outside of the structures; external spots appeared between cells, at cleft-like sites. Migration-permissive hydrogels (PQ-RGD) supported structures with similar laminin expression patterns, but with a higher proportion of external laminin deposits. Moreover, the signal intensity for those external laminin expression spots was higher and more condensed then for those in the migration-impermissive hydrogels. Fibronectin expressed by structures in HA-PEGDA appeared inside and outside, with most of fibronectin appearing to be localized within the cells (Figure 7b). In peptide-modified hydrogels (PQ-RGD and PQC-RDG), fibronectin was localized to external “cleft-like” invaginations on the periphery of each structure (Figure 7b).

**Figure 7.**
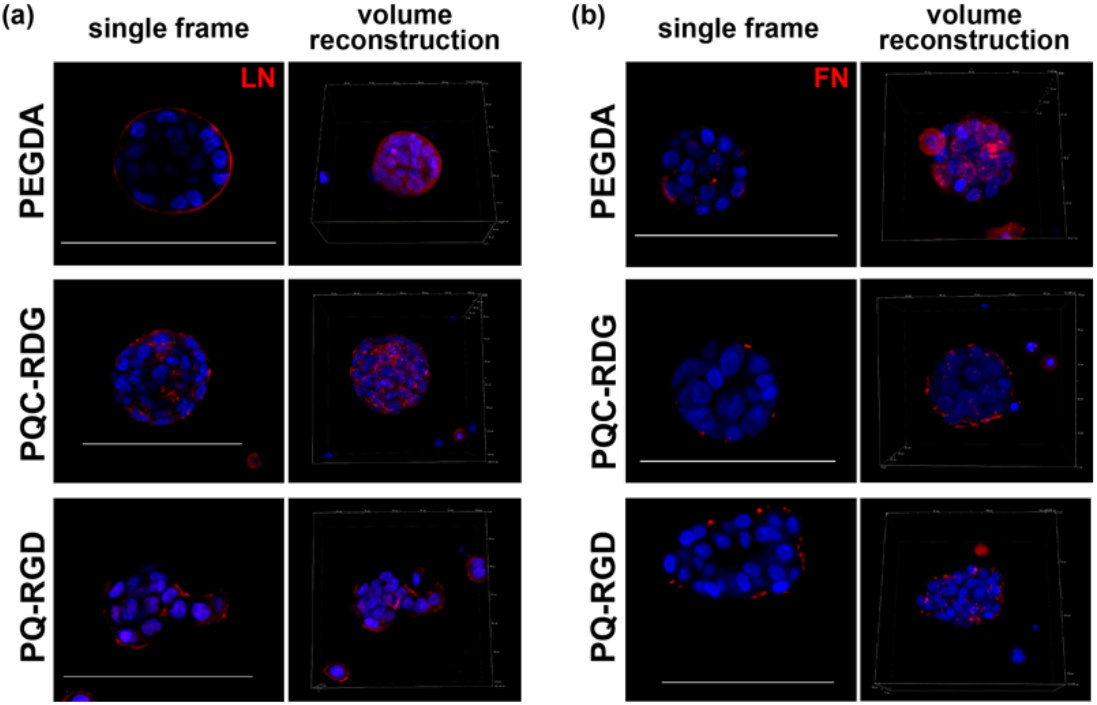
Extracellular matrix deposition. Panels show representative single frame images and 3D Z-stack volume reconstructions of immunofluorescence staining (in red) for the ECM proteins (a) I laminin (LN) and (b) fibronectin (FN) in hS/PC-laden hydrogel variants after 14 days of culture. hS/PCs in HA-PEGDA hydrogels localized both ECM proteins to the spheroid peripheries, with LN particularly observed as a thick band of basement membrane. Conversely, hS/PCs in peptide-modified hydrogels localized LN and FN to interstitia between cells at invaginations. Nuclei are labeled blue. n ≥ 3 for all conditions. Scale bars = 100μm.

To investigate if the fibronectin deposition pattern was consistent with early clefting of glands, hS/PC structures were immunostained concurrently for fibronectin, fibronectin receptor integrin α_5_, and branching morphogenesis associated protein Btbd7 at day 14 (Figure S7b). Fibronectin expression was consistent with the previous description. However, integrin α_5_ expression was lost in migration-permissive hydrogels by day 14. Interestingly, Btbd7 expression was highest in hS/PC structures in migration-permissive hydrogels, with most of its expression being in cells located towards the outer structure and next to fibronectin deposits.

### 3.4. Phenotype of hS/PC structures

To evaluate if hS/PC interaction with the distinct surrounding matrices promoted differentiation, hS/PCs encapsulated in 11:1 (SH:Ac) hydrogels were immunostained with various phenotypic markers (Figure 8). Broad expression of Keratin 5 (K5) and K14 confirmed retention of progenitor markers. Myoepithelial marker α-smooth muscle actin (ACTA2) was never observed, and ductal marker Keratin 19 (K19) was very rarely observed in the hS/PC cultures. Acinar marker α-amylase was present at low levels throughout hS/PC structures, consistent with prior observations of hS/PCs.

**Figure 8.**
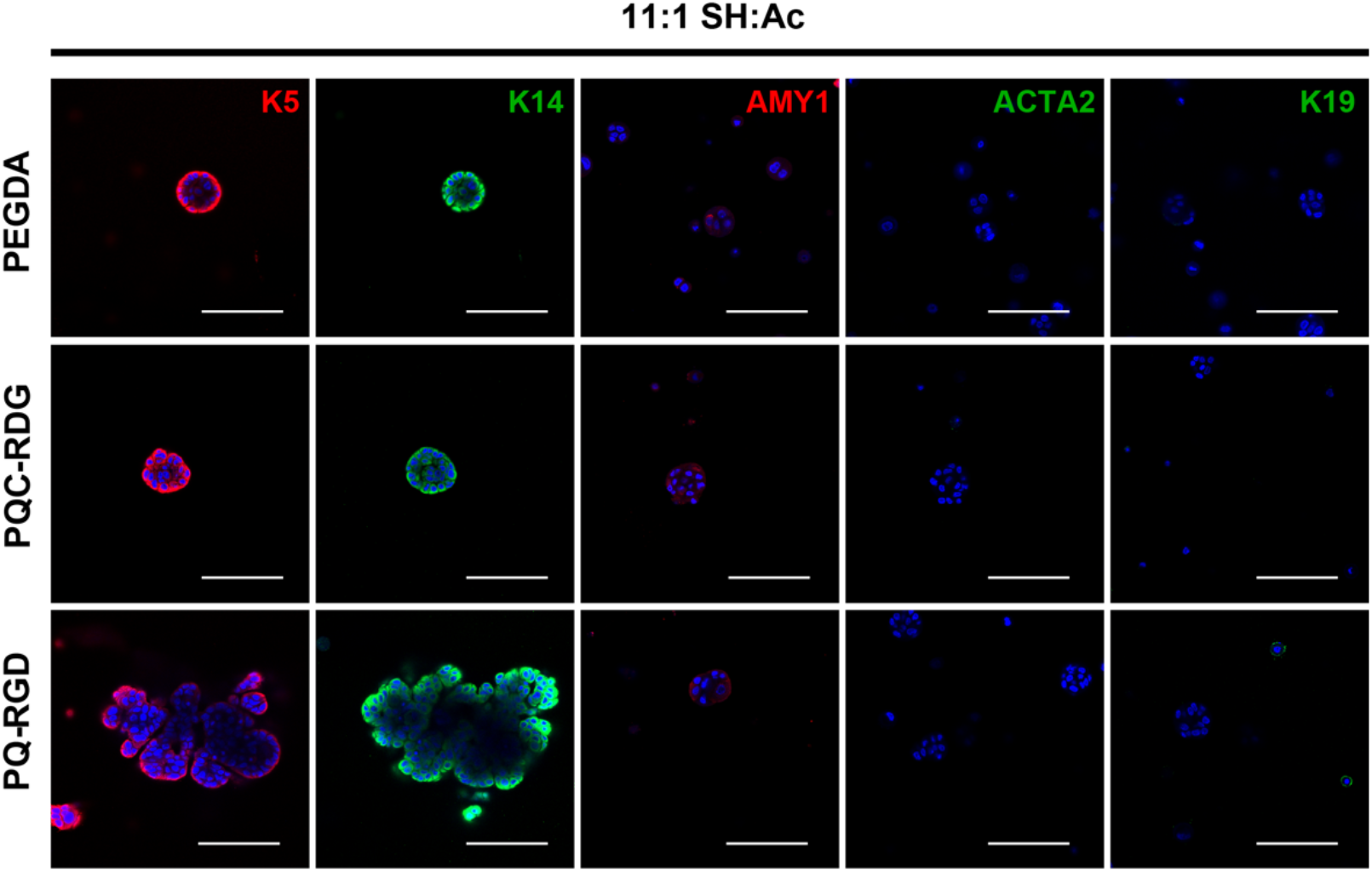
Assessment for hS/PC differentiation. Representative single-frame images of hS/PCs immunostained for various phenotypic markers at day 14. hS/PCs retained expression of progenitor markers Keratin 5 (K5; red) and K14 (green) and low levels of acinar marker a-amylase (AMY1; red). No hS/PCs were observed to express myoepithelial marker a-smooth muscle actin (ACTA2), and only rare hS/PCs expressed detectable levels of ductal marker K19.

## 4. Discussion

The extensive damage caused to the SGs after radiation therapy predominantly impacts acinar cells, and their antecedent progenitor cell source. Our lab proposes to generate a functional, tissue-engineered SG replacement to restore salivary function in patients suffering from xerostomia. Creating a biocompatible salivary gland substitute will require a matrix that supports the self-organization of the encapsulated therapeutic cells (hS/PCs and others), as well as integration with patient tissue surrounding the implant.

Although base HA-PEGDA hydrogels have been instrumental for baseline assessments of hS/PC characteristics such as growth and phenotype [26,27,36,37], these matrices have some limitations for use in bioengineering salivary gland constructs that resemble the native glandular structure. At early stages of SG formation, branching morphogenesis is a key step that establishes the gland’s characteristic branched structure and directional secretion. Characteristic of this process is the deposition of punctate FN at the cleft points between branching cell buds. These phenotypic behaviors of peripheral clefting and punctate FN deposition are rarely observed for hS/PCs cultures in HA-PEGDA matrices. Fortunately, the modular chemistry of these hydrogels enables the covalent addition of biofunctional peptides that expand the hydrogels’ capacity to support cell migration and more complex organization. Integration of the “PQ” and RGD moieties creates a migration-permissive hydrogel (MP-HA) that yields the behaviors observed here for hS/PCs, will enable future co-cultures with other stromal cell types, and supports biodegradation of the matrix, as each is required for complete implant integration into the patients’ tissue bed.

Because of the introduction of these modifying groups, all of the peptide-modified hydrogels (PEGDA-RGD, PQC-RDG, PQC-RGD, PQ-RGD, PQ-RDG), at all crosslinking densities, were physically larger (i.e. a higher swelling ratio) than their HA-PEGDA counterparts by day 1 (Figure S5). Notably, all of the pendant RGD-RDG peptides were covalently attached to the gels using a PEG-Ac tether, and all of the bifunctional peptide-based PQ-PQC crosslinkers were constructed with dual PEG-Ac chains, one on either side. Therefore, all of the peptide-modified hydrogels contained proportionally higher PEG content than the base HA-PEGDA hydrogels, which can lead to a higher gel swelling ratio.

Additionally, as noted in Figure 1a, the base HA-PEGDA hydrogels crosslink in a two-step process: acrylates first consume thiols quickly to form initial covalent bonds, and then residual thiols further react over several hours to form additional disulfide crosslinks [38]. Our peptide-modified variants consume many of these thiols, through attachment of the acrylated pendant integrin ligands, leading to a reduced level of secondary disulfide crosslinks. Therefore, the combination of these two effects (higher total PEG content and fewer disulfide-based crosslinks) would be expected to produce gels with higher networklevel porosity, and larger macroscopic size, as was seen in Figure S5. Prior studies of similar systems demonstrated an extended impact of residual disulfide formation over time, and we observe similar impacts in this work, particularly in the contraction of HA-PEGDA gels after initial formation. Moreover, we performed rheological characterization of the hydrogels used in these studies because we were interested in whether these varying factors would have an effect on the hydrogel viscoelastic properties. We observed that peptide-modified hydrogels had a consistently higher storage modulus (G’) than the base HA-PEGDA system at each crosslink density, again likely due to the enhanced swelling from PEG content.

This work showed that hydrogel crosslink density, swelling and biofunctional peptide incorporation influenced hS/PCs multicellular structure viability, size, structure formation frequency, and morphology (Figures 2, 3). Cell viability in base HA-PEGDA hydrogels was only significantly lower than that in peptide-modified hydrogels at 6:1 SH:Ac ratios; otherwise, cell viability was comparable among all hydrogels at 11:1 and 16:1. Peptide-modified hydrogels, regardless of the peptides integrated, led to larger multicellular structures; thus, it seems that increased pore size in hydrogels with a higher swelling ratio is an important factor affecting hS/PC structure size, and potentially mobility. Integration of RGD in hydrogels with either a noncleavable or cleavable crosslinker increased the number of multicellular structures by day 7. Moreover, substitution of PQC with PQ in conditions lacking RGD had no effect on multicellular structure formation over time. Therefore, having an integrin-binding site in the hydrogel matrix was more important than having an MMP cleavable crosslinker to increase the number of hS/PC structures formed. It is important to recognize that these studies were performed with only one MMP-degradable crosslinker; there is clear relevance to studying a panel of substrate specific preferences by hS/PCs, or by other associated salivary fibroblasts, endothelium, nerve, etc. In future clinical use, integrating both adhesive and degradable peptides into the matrix during hydrogel design would enable further customization: RGD incorporation into the encapsulation matrix would shorten the time that hS/PCs need to be cultured to form multicellular structures before implantation and PQ incorporation into the encapsulation matrix would enable biodegradation needed for proper implant integration.

Interestingly, the percentage of proliferating cells was significantly higher in MP-HA than in other hydrogel variants crosslinked at 6:1 and 16:1. When we systematically incorporated biofunctional peptides, we observed that integrating RGD into hydrogels crosslinked with either of the three crosslinkers (PEGDA, PQC, and PQ) always led to significantly increased proliferation by hS/PCs. However, hS/PC proliferation was unaffected in hydrogels with pendant RDG crosslinked with cleavable PQ when compared to those crosslinked with PEGDA or PQC. Taken together, this demonstrates that presence of integrin ligands in the scaffold is most important for stimulating hS/PCs to proliferate. However, the enzymatically-degradable peptide will be essential for future studies that will also include a stromal salivary compartment, which will be crucial for the regeneration of salivary glands in a damaged tissue bed.

Developing salivary glands and their surrounding mesenchyme secrete ECM proteins, including fibronectin, to signal salivary cells to grow, proliferate, and differentiate into their adult phenotypes (acinar, ductal and myoepithelial) [39,40]. This signaling is mediated by integrin receptors that bind to ECM ligands such as RGD and activate intracellular signaling pathways needed for cell survival, proliferation, and differentiation [41,42]. Integrin β_1_ is commonly expressed throughout the various stages of salivary gland development, and its expression continues in the differentiated adult gland [43,44]. We showed that hS/PCs in MP-HA always demonstrated increased activated integrin β_1_ over the other hydrogel variants at all crosslink densities. In our systematic approach of adding individual biopeptides to assess their role on integrin β_1_ activation (11:1; SH:Ac), we discovered that integrating RGD in hydrogels crosslinked with either of three crosslinkers consistently led to increased activation of integrin β_1_, when compared to their respective controls. Additionally, we showed that increased internal 3D space created by larger crosslinkers/PEG content independent of RGD promoted increased activation of integrin β_1_. Still, concurrent integration of PQ and RGD had the greatest impact on integrin β_1_ activation. We further confirmed that MP-HA supports the most cell-ECM interactions by identifying higher expression levels of fibronectin and laminin integrin receptors in this system. Thus, migration-permissive hydrogels offer a significant advance over base HA-PEGDA hydrogels, because they promote enhanced integrin β_1_ activation by hS/PCs, which is key for guiding hS/PCs to grow, organize and differentiate into a functional salivary gland.

Space for cell growth and organization provided by the tissue engineering scaffold is also central for bioengineering a branched glandular structure. hS/PCs encapsulated within peptide-modified hydrogels formed multicellular structures with cleft-like sites around the structures’ peripheries by day 14. Particularly, MP-HA promoted formation of larger, multicellular structures with morphologies, i.e., presence of clefts and lumen-like cavities, similar to a developing salivary gland (initial bud, pseudoglandular, and canalicular stages). However, cells in base hydrogels formed tight spherical structures that may not be able to re-organize into larger glandular (branched) structures.

Lastly, we discovered that basement membrane deposition patterns parallel the changes in hS/PC structure morphology induced by peptide-modified hydrogels. Basement membrane deposition by hS/PCs in migration-permissive hydrogels resembles that of a developing salivary gland; it is localized at cleft sites allowing protrusions/buds of cells to grow outward. Additionally, the expression pattern of branching morphogenesis associated protein Btbd7 by hS/PC structures in MP-HA was similar to that *in vivo* during gland morphogenesis, which further strengthened the conclusion that MP-HA supports the organization of hS/PCs into glandular-like structures [45]. Assessment of hS/PCs for differentiation markers in MP-HA via immunostaining experiments showed retention of a progenitor phenotype (K5^+^, K14^+^; Figure 8), and no other clear markers of differentiation, which suggests that these hS/PC structures have not reached the later stages of development (terminal differentiation). In contrast, hS/PCs in HA-PEGDA localize basement membrane evenly around the structures’ peripheries, appearing to be restrictive of growth. Therefore, MP-HA should better enable cell growth and branching than undecorated counterparts. Nevertheless, other aspects important for salivary gland development, such as instructive cell types neighboring the gland (neuronal, endothelial, mesenchymal, etc.) and additional extracellular matrix components, would further improve this tissue construct model, expectantly fulfilling requirements needed for hS/PC differentiation into a functional, mature salivary gland.

Migration-permissive MP-HA hydrogels meet the basic requirements for integration of the implant to the patient’s surrounding tissue. Specifically, MP-HA contains enzymatically degradable crosslinkers to enable timely angiogenesis and creation of larger pores, or spaces, within the matrix and fibronectin-derived integrin-binding sites to support cell adhesion needed for cell migration, and ultimately, organization of hS/PCs into salivary-like structures. Although hydrogel matrices with PQ-RGD have been employed by many laboratories for a variety of uses, such features have not been applied to SG regeneration using a systematic approach as shown by this work. PEG-based matrices, degradable either by MMPs or hydrolysis, have been used previously for culture of preformed multicellular spheroids isolated from mouse submandibular glands [46]. However, these matrices did not include RGD ligands to target integrin receptors on the mouse-derived cells. Conversely, HA-SH matrices have been described with RGD functionalization, via different chemistries, for ocular engineering applications [47], but these did not include PQ crosslinkers nor as much PEG, both of which were observed in the present work to greatly influence hS/PC organization into morphologies and ECM patterns that are closer to that of the native SG. In both of these example papers, the degree of hydrogel porosity is difficult to compare to that in the present work, however it is clear that open space within a hydrogel network greatly impacts key characteristics. Of course, the protein-based murine system Matrigel^®^ supports branching, but cannot be used *in vivo* in humans.

Instead, we modify HA-based substrates with the minimum necessary peptide features to enable cell-ECM interplay. Importantly, MP-HA matrices enable the incorporation of integrin-dependent salivary-derived stroma, which can participate in a coculture model. This is a topic for future work from our laboratory.

## 5. Conclusions

MP-HA hydrogels are an improved hydrogel support matrix for bioengineering glandular organs, such as salivary glands, and expand the options for cell-ECM dialog required for development. Integration of an integrin-activating peptide and greater internal 3D space yields high cell viability, promotes formation of large multicellular structures, leads to a higher number of multicellular structures, and promotes increased cell proliferation—all of which are important attributes in a successful tissue engineering model. Moreover, peptide-modified hydrogels enable hS/PCs to organize into asymmetric multicellular structures with punctate basement membrane deposition patterns characteristic of a developing, branched SG. Concurrent incorporation of cleavable PQ and RGD in MP-HA significantly enhances activation of integrin β_1_ shortly after encapsulation. Furthermore, MP-HA meets the basic requirements of a tissue engineering scaffold conducive to generating a glandular tissue construct that could support integration of vasculature, innervation and mesenchymal cells from the patient for long term relief from xerostomia.

## Acknowledgments

This work was funded by NIH R03DE028988 to DAH, NIH R01DE022969 to MCFC, NSF GRFP DGE-1450681 to MM, NSF IRISE 0966303 to MM, UTHealth Startup Funds to DAH, Rice University AGEP / Office of Diversity Inclusion to MM, and philanthropic donations. Instruments used in this work were supported by a UT STARS award to MCFC. We would like to acknowledge Eliza L. S. Fong and the rest of the salivary team (Danielle Wu, Kelsea Hubka) for many helpful discussions.

## Data Availability

Raw data primarily comprise confocal microscopy images and are retained for a sufficient time in accordance with NIH data retention guidelines. These are available on an individual basis upon reasonable request. All processed images are presented in the manuscript.

---

**Supplementary Table S1.** Information for antibodies used in this study.

| Immunogen | Host | Class | Company | Catalog # | Stock Concentration | Working Dilution |
| --- | --- | --- | --- | --- | --- | --- |
| **Primary Antibodies** |  |  |  |  |  |  |
| Btbd7 | rabbit | polyclonal | Atlas Antibodies<br>Stockholm,<br>Sweden | HPA049926 | 0.1 mg/mL | 1:100 |
| Fibronectin (FN) | rabbit | polyclonal | abcam,<br>Cambridge, MA | ab2413 | 0.14 µg/mL | 1:100 |
| Integrin alpha 1<br>(ITGA1) | mouse | monoclonal | Novus<br>Biologicals,<br>Centennial, CO | NBP2-29757 | 1 mg/mL | 1:100 |
| Integrin alpha 5<br>(ITGA5) | mouse | monoclonal | Novus<br>Biologicals,<br>Centennial, CO | NBP2-50146 | 1 mg/mL | 1:100 |
| Integrin alpha 5<br>(ITGA5) | rabbit | monoclonal | abcam,<br>Cambridge, MA | ab150361 | 0.22 mg/mL | 1:100 |
| Integrin beta 1<br>(ITGB1) | mouse | monoclonal | Novus<br>Biologicals,<br>Centennial, CO | NB100-63255 | 1 mg/mL | 1:100 |
| Integrin beta 4<br>(ITGB4) | mouse | monoclonal | abcam,<br>Cambridge, MA | ab29042 | N/A | 1:100 |
| Pan-Laminin | rabbit | polyclonal | Invitrogen,<br>Carlsbad, CA | PA1-16730 | 1 mg/mL | 1:100 |
| Ki67 (MKI67) | mouse | monoclonal | Santa Cruz<br>Biotechnology,<br>Dallas, TX | SC-101861 | N/A | 1:100 |
| **Secondary Antibodies** |  |  |  |  |  |  |
| mouse IgG<br>(AlexaFluor 488-linked) | goat | polyclonal | Invitrogen,<br>Carlsbad, CA | A11029 | 2 mg/mL | 1:500 |
| rabbit IgG<br>(AlexaFluor 568-linked) | goat | polyclonal | Invitrogen,<br>Carlsbad, CA | A11011 | 2 mg/mL | 1:500 |

**Supplementary Table S2.** Information for counterstains and cell dyes used in this study.

| Cell dye/<br>Counterstain | Company | Catalog # | Stock Concentration | Working Concentration |
| --- | --- | --- | --- | --- |
| calcein AM | Biotium,<br>Hayward, CA | 80011-3 | 4 mM | 2 µM |
| DAPI | Thermo Fisher<br>Scientific, Grand<br>Island, NY | D1306 | 5 mg/mL | 2 µg/mL |
| ethidium<br>Homodimer III | Biotium,<br>Hayward, CA | 40050 | 2 mM | 4 µM |
| Hoechst 33342 | Enzo Life<br>Sciences, Inc.,<br>Farmingdale, NY | ENZ-52401 | 10 mg/mL | 10 µg/mL |
| Alexa Fluor™<br>633 Phalloidin | Invitrogen,<br>Carlsbad, CA | A22284 | 6.6 µM | 0.165 µM |

**Supplementary Table S3.** Concentrations of working solutions of crosslinkers (CL) and pendant peptides (PP) used at a 4:1:1 (thiolated HA:CL:PP) volume ratio to make 42 μL hydrogels. Final concentrations of CLs used in hydrogels with crosslinking densities of 6:1, 11:1 or 16:1 (SH:Ac) were 0.46, 0.25 or 0.17 mM. Final concentration of PPs used in peptide-modified hydrogels was 3 mM.

|  | Crosslinking Density |  |  |  |  |  |
| --- | --- | --- | --- | --- | --- | --- |
|  | 6:1 (SH:Ac) |  | 11:1 (SH:Ac) |  | 16:1 (SH:Ac) |  |
| Component | CL<br>(2.77 mM) | PP (<br>18 mM) | CL<br>(1.51 mM) | PP<br>(18 mM) | CL<br>(1.04 mM) | PP<br>(18 mM) |
| PEGDA<br>(mg/mL) | 9.42 | --- | 5.14 | --- | 3.53 | --- |
| PEGDA/RGD<br>(mg/mL) | 9.42 | 70.0 | 5.14 | 70.0 | 3.53 | 70.0 |
| PQC/RDG<br>(mg/mL) | 22.3 | 70.0 | 12.2 | 70.0 | 8.35 | 70.0 |
| PQC/RGD<br>(mg/mL) | 22.3 | 70.0 | 12.2 | 70.0 | 8.35 | 70.0 |
| PQ/RDG<br>(mg/mL) | 22.5 | 70.0 | 12.3 | 70.0 | 8.43 | 70.0 |
| PQ/RGD<br>(mg/mL) | 22.5 | 70.0 | 12.3 | 70.0 | 8.43 | 70.0 |

## Notes

**Conflict of Interest:** None.

### Competing Interest Statement

The authors have declared no competing interest.

